# The kinases PIG-1 and PAR-1 act in redundant pathways to regulate asymmetric division in the EMS blastomere of *C. elegans*

**DOI:** 10.1101/191866

**Authors:** Malgorzata J. Liro, Diane G. Morton, Lesilee S. Rose

## Abstract

The PAR-1 kinase of *C. elegans* is localized to the posterior of the one-cell embryo and its mutations affect asymmetric spindle placement and partitioning of cytoplasmic components in the first cell cycle. However, unlike mutations in the posteriorly localized PAR-2 protein, *par-1* mutations do not cause failure to restrict the anterior PAR polarity complex. Further, it has been difficult to examine the role of PAR-1 in subsequent divisions due to the early defects in *par-1* mutant embryos. Here we show that the PIG-1 kinase acts redundantly with PAR-1 to restrict the anterior PAR-3 protein for polarity maintenance in the one-cell embryo. By using a weak allele of *par-1* that exhibits enhanced lethality when combined with a *pig-1* mutation we have further explored roles for these genes in subsequent divisions. We find that both PIG-1 and PAR-1 regulate spindle orientation in the EMS blastomere of the four-cell stage embryo to ensure that it undergoes an asymmetric division. In this cell, PIG-1 and PAR-1 act in parallel pathways for spindle positioning, PIG-1 in the MES-1/SRC-1 pathway and PAR-1 in the Wnt pathway.

## Introduction

Asymmetric divisions are important for cell fate diversity during development and for stem cell maintenance. In the process of asymmetric cell division, cell fate determinants become asymmetrically distributed along an axis of polarity. Concurrently, the mitotic spindle aligns along the same polarity axis and signals the cleavage plane to bisect the determinant asymmetry, ultimately producing two differentially fated daughter cells. Incorrect distribution of cell fate determinants as a result of mitotic spindle misorientation has been linked to changes in cellular proliferation, cell fate specification, and cancer (Bergstralh et al., 2017; Neumuller and Knoblich, 2009).

The *Caenorhabditis elegans* one-cell embryo is a classical model system for studying the process of asymmetric cell division. The sperm entry point marks the posterior end of the embryo and the sperm centrosomes are required for initiating symmetry breaking, which then leads to the establishment of anterior/posterior polarity. Symmetry breaking is characterized by the activation of cortical contractility; a cortical acto-myosin network, regulated by the non-muscle myosin II heavy chain NMY-2, flows away from sperm and associated centrosomes. The posterior, non-contractile cortex expands towards the anterior until it reaches about 50% embryo length, and a prominent pseudocleavage furrow is present at the border of the contractile and non-contractile domains. These contractility differences along the one-cell cortex occur concurrently with the asymmetric segregation of PAR proteins.

PAR polarity proteins were named after their mutant phenotype; failure of asymmetric *par*titioning of cytoplasmic components, including cell fate determinants in the one-cell embryo, results in altered cell fate, cell size, cell cycle timing, and often abnormal spindle positioning, which leads to catastrophic defects in developmental patterning (Kemphues et al., 1988). Several of the PAR polarity proteins are polarized along the anterior/posterior (AP) axis during polarity establishment. The PDZ-containing proteins PAR-3 and PAR-6 and the atypical protein kinase C, PKC-3, initially localize uniformly around the entire cell cortex, then become localized to the anterior domain, allowing PAR-2, a RING-finger protein, and PAR-1, a Ser/Thr kinase, localize to the reciprocal posterior domain of the embryo (Goldstein and Macara, 2007; Rose and Gonczy, 2014; Wu and Griffin, 2017). Following their establishment in these domains, PAR proteins mutually exclude each other from their distinct anterior and posterior domains to maintain cell polarity. PAR proteins are required for posterior localization of the cell fate determinant PIE-1 (Tenenhaus et al., 1998) and germline P granules (Kemphues et al., 1988) as well as for anterior segregation of somatic precursor cell fate determinants such as MEX-5 (Schubert et al., 2000). The distinct PAR domains are also important for regulating other cortical proteins that control the alignment of spindle with cell fate determinants to ensure generation of properly fated daughter cells following cell division (di Pietro et al., 2016; Rose and Gonczy, 2014).

The *par-1* gene was initially identified by maternal effect lethal mutations that disrupt asymmetric spindle placement and partitioning of cytoplasmic components in the one-cell embryo (Kemphues et al., 1988). *par-1* mutants have more symmetric first cleavage and synchronous cell cycles at the two-cell stage, similar to that observed in *par-2* mutants. At the same time, the effect of *par-1* mutations on maintenance of the PAR polarity domains and spindle orientation at the two-cell stage is much weaker than that of PAR-2, raising the possibility that another protein functions redundantly with PAR-1 in these processes (Cuenca et al., 2003; Etemad-Moghadam et al., 1995; Kemphues et al., 1988).

Morton and colleagues identified a set of genes that when knocked down by RNA interference (RNAi) do not cause high embryo lethality on their own, but cause higher frequency of embryo lethality when combined with the *par-1(zu310ts)* mutation at semi-permissive temperature (Morton et al., 2012). One of these genes was *pig-1*, which encodes a serine/threonine kinase related to the PAR-1 kinase and orthologous to vertebrate Maternal Embryonic Leucine zipper Kinase (MELK). PIG-1 was originally described for its role in regulating cell size and cell fate during asymmetric division of neuroblasts in the late embryo and L1 larva (Cordes et al., 2006). *pig-1(RNAi)* also enhanced the lethality of *par-2(ts*) embryos at semi-permissive temperature and *pig-1(RNAi); par-2* embryos showed greater defects in maintaining the anterior PAR domain boundary in the one-cell embryo and stronger defects in two-cell asymmetries compared to *par-2(ts)* alone (Morton et al., 2012). These observations indicate that PIG-1 plays a role in early polarity, but neither the *pig-1* early embryonic phenotype nor its potential redundancy with *par-1* in the one-cell embryo have been analyzed in detail.

PAR domains are re-established in the posterior P1 cell after the first division, and P1 then divides asymmetrically to produce another germ-line precursor, P2, and the endo-mesodermal precursor cell, EMS. The EMS cell divides asymmetrically to produce the anterior MS daughter cell, which primarily produces mesodermal cells, and the smaller posterior daughter cell, E, which gives rise to the entire endoderm of the worm. The EMS division orientation is due to a 90° rotation of the nuclear-centrosome complex that occurs prior to nuclear envelope breakdown (NEB) such that spindle forms on the cell’s A/P axis. Blastomere isolation and reconstitution experiments showed that both the division orientation and the fate asymmetry of the EMS daughter cells depend on a signal from the P2 cell (Goldstein, 1993, 1995a, b; Maduro, 2017; Rose and Gonczy, 2014). Subsequent genetic screens for mutants producing more mesoderm at the expense of endoderm tissue (“mom” mutants) identified several components of the Wnt signaling pathway as being partially required for endoderm specification. A subset of Wnt signaling components, such as the Wnt ligand (MOM-2 in C. elegans), the Frizzled receptor (MOM-5), and the Disheveled adaptor proteins (DSH-2 and MIG-5) are also partially required for timely spindle orientation. Embryos mutant for these components can exhibit a failure of nuclear rotation by NEB, but most embryos eventually orient their spindles onto the A/P axis in anaphase (Bei et al., 2002; Liro and Rose, 2016; Schlesinger et al., 1999).

A similar late spindle rotation phenotype in the EMS cell is exhibited by mutants in the MES-1/SRC-1 pathway (Bei et al., 2002). However, when embryos are mutant for components in both the Wnt and MES-1/SRC-1 pathways, EMS spindle positioning fails, resulting in divisions along the left/right (L/R) axis in nearly all embryos (Bei et al., 2002). Although *mes-1* and *src-1* single mutants do not have defects in endoderm fate specification, double mutants with Wnt pathway components produce a higher frequency of failure to specify endoderm than in *wnt* single mutants. Thus, the MES-1/SRC-1 pathway acts in parallel to the MOM-2/Wnt pathway for both EMS spindle orientation and intestinal specification (Bei et al., 2002).

The MES-1/SRC-1 pathway has only a few known components. SRC-1, a tyrosine kinase, and MES-1, a transmembrane receptor tyrosine kinase-like protein, both localize to the cell-cell interface between P2 and EMS (Bei et al., 2002; Berkowitz and Strome, 2000). NMY-2 was shown to be upstream of SRC-1 in intestinal specification, but not spindle positioning (Liu et al., 2010). Recently, LET-99, a protein required for spindle positioning in the one-cell embryo (Krueger et al., 2010; Park and Rose, 2008; Rose and Kemphues, 1998; Tsou et al., 2002; Wu and Rose, 2007), has been shown to play a role in EMS spindle positioning, where it appears to act downstream of the MES-1/SRC-1 signal (Bei et al., 2002; Liro and Rose, 2016). In the one-cell embryo, LET-99 is asymmetrically localized by the PAR proteins at the cortex, where it then acts to localize the force generation complex that aligns the spindle on the PAR polarity axis.

However, in the EMS cell, the PAR domains localize to distinct inner and outer domains (Nance and Priess, 2002), and thus are not aligned with the A/P axis and the spindle as they are in the one-cell. Whether *par* genes are required during EMS asymmetric division has not been assessed, due to their mutant phenotypes prior to this stage.

Here, we further investigate the roles of *pig-1* in the *C. elegans* embryo development and how *pig-1* may be synergizing with *par-1*. We find that *pig-1* acts in parallel with *par-1* in regulating restriction of the anterior polarity complex at the one-cell stage. We further find that PIG-1 acts to regulate EMS spindle orientation in the MES-1/SRC-1 pathway and PAR-1 in the parallel Wnt pathway. Moreover, in addition to its role in spindle positioning, PIG-1 appears to play a role in endoderm cell fate specification.

## Results

### PIG-1 acts with PAR-1 to restrict the anterior PAR domain in the one-cell *C. elegans* embryo

While the *pig-1(gm344)* null mutant strain (Cordes et al., 2006) is generally homozygous viable, approximately 9% of embryos fail to hatch, and approximately 15% of larvae die before reaching adulthood at 19.5°C (Table S1). Thus, the *pig-1* gene, while not essential for viability, makes a clearly discernable contribution to development. We compared viability at different temperatures, and found that at a higher temperature of 25°C, embryo lethality was increased to 14 % in *pig-1(gm344)* mutants (hereafter referred to as *pig-1* mutants; Table S1). The terminal phenotypes of the dead embryos showed that they had many differentiated cell types, but lacked morphogenesis. This phenotype has also been observed in embryos produced by mothers homozygous for maternal effect lethal mutations in the *par* genes (Kemphues et al., 1988). In addition, *pig-1(RNAi)* has been found to enhance the lethality of *par-1* and *par-2* temperature-sensitive mutants (Morton et al., 2012). We therefore examined early embryogenesis of the *pig-1* mutant to determine whether there were any defects in early polarity, using time-lapse microscopy.

Before pronuclear meeting in the newly-fertilized egg, a series of myosin-driven contractions facilitates a rearrangement of the cortex and allows polarization of cortical PAR proteins into their anterior and posterior domains. A signature of this reorganization is the presence of anterior cortical contractions that resolve into a single pseudocleavage constriction at mid-embryo length, which then relaxes after pronuclear meeting. Time-lapse imaging revealed that *pig-1* embryos grown at either 19.5°C or 25°C exhibited less anterior cortical ruffling and had a diminished pseudocleavage furrow (N= 10 embryos at each temperature), as compared to wild type (N= 12) (Figure 1A, Movie S1, S2). Nonetheless, the first cleavage spindles were aligned with the A/P axis and displaced towards the posterior in *pig-1* embryos, such that the two daughter cells had different sizes, as in wild type.

**Figure 1.**
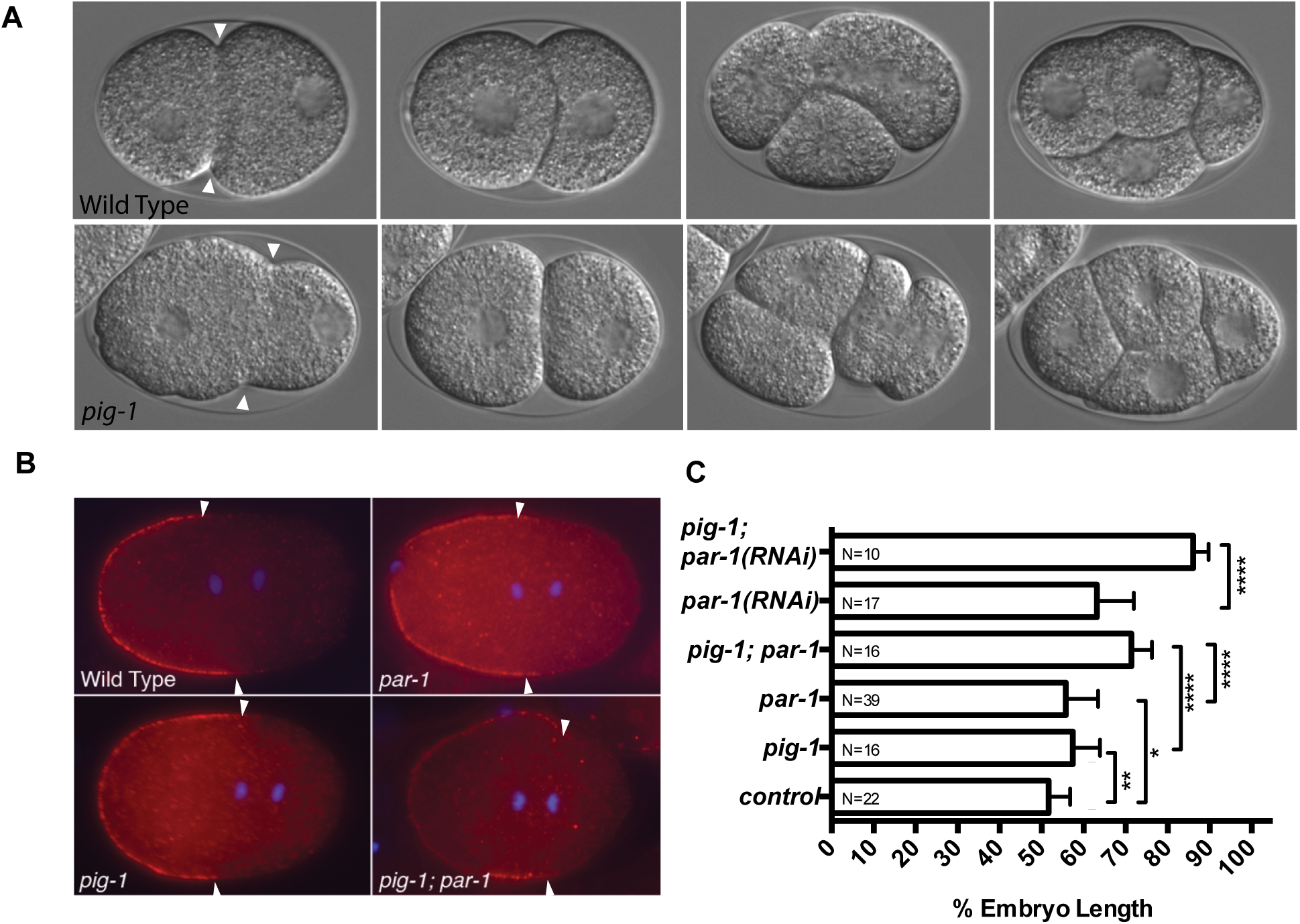
*pig-1* mutants exhibit reduced contractility and expansion of the PAR-3 domain. (A) Images from time-lapse microscopy of wild-type and pig-1(gm344) embryos grown and imaged at 19.5°C. Anterior is to the left and posterior to the right in this and all embryo images. Arrowheads mark the pseudocleavage furrow, which is reduced in pig-1. (B) Representative images of one-cell embryos at anaphase showing anti-PAR-3 antibody staining (red) in wild-type control (N2), *pig-1(gm344)*, *par-1(zu310ts)* or *par-1(RNAi)* single and double mutants grown at 25°C. Arrowheads mark the extent of the PAR-3 domain. DAPI (blue) marks the DNA. (C) Quantification of the cortical PAR-3 signal expressed as percent embryo length, from anterior 0% to posterior 100%. Data were compared using the unpaired Student’s t-test (* P ≤ 0.05; **P ≤ 0.01; ****P ≤ 0.0001; see Table S2 for all P values).

Wild-type embryos also exhibit characteristic differences in cleavage patterns at second and third cleavage that are dependent on normal PAR function (Figure 1A). At second cleavage, the anterior AB cell divides before the P1 cell (126 sec on average, Figure 2A); the P1 nucleus exhibits a nuclear rotation event so that the spindle forms on the A/P axis once again. At third cleavage, the anterior EMS daughter divides before the P2 cell (232 sec on average), with its spindle also orientated on the A/P axis. Examination of *pig-1* mutant embryos revealed that the AB and P1 daughter cells divided with more similar cell cycle rates than in wild type (96 sec; Figure 2A). Nonetheless, the P1 nucleus rotated on to the A/P axis in *pig-1* mutants, just as in wild type (N = 10). At the third division, the difference between the EMS and P2 cell cycles of *pig-1* mutants (214 sec) was not statistically different from wild type, and most embryos showed normal EMS division patterns (Figure 2A,B and see below). Overall, the phenotypes exhibited for *pig-1* embryos are less severe than those observed for most *par* mutant embryos, but nevertheless show that PIG-1 is important for some of the normal asymmetries of the early blastomeres.

**Figure 2.**
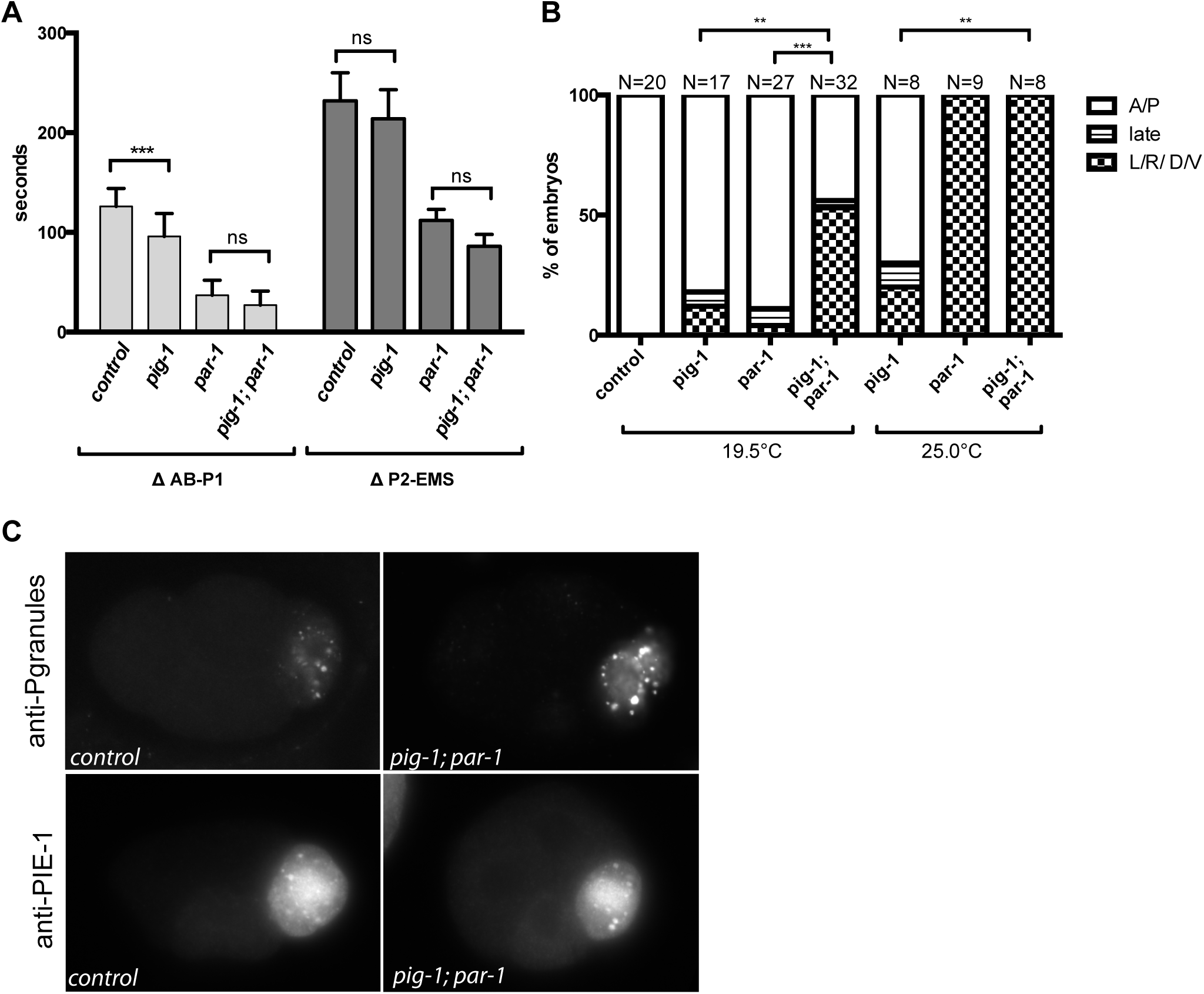
Comparison of *pig-1* to *pig-1;par-1* double mutant embryos. (A) Cell cycle comparison between wild-type controls, *pig-1(gm344), par-1(zu310ts),* and double mutant embryos. The difference between the time of onset of nuclear envelope breakdown for AB-P1 and EMS-P2 sister cells is shown on the y-axis. Data were compared using the unpaired Student’s t-test. (B) Quantification of spindle positioning in *pig-1(gm344), par-1(zu310)* and double mutants at the temperatures indicated. The A/P category includes embryos whose centrosomes were aligned on the A/P axis before nuclear envelope breakdown. The late rotation category refers to EMS spindle alignment with the A/P axis that occurred after NEB; the L/R / D/V category includes final spindle positions on all non-A/P axes. The proportion of normal (A/P) and abnormal (late and L/R D/V) embryos were compared between genotypes using Chi-squared analysis. (ns, not significant, P>0.05; * P ≤ 0.05; **P≤0.01; *** P ≤ 0.001; see Table S2 for specific P values). (C) Representative images of P-granule and PIE-1 localization in control and *pig-1(gm344);par-1(zu310)* double mutant embryos.

The abnormalities in cortical contractility exhibited by *pig-1* embryos suggested that there might also be defects in PAR domain establishment or maintenance. Previous work showed that although *par-1* mutant zygotes have a distinct anterior PAR-3 domain, the extent of the domain expands slightly towards the posterior during the maintenance phase in *par-1(RNAi*) one-cell embryos (Cuenca et al., 2003); these observations suggest that PAR-1 may participate in restriction of the anterior polarity complex, but perhaps redundantly with another protein. Because of the structural similarities between PIG-1 and PAR-1 kinases as well as their synergy in causing embryo lethality when the activities of both are reduced (Morton et al., 2012)(Table S1), we wanted to determine how *pig-1* and *par-1* together might affect PAR polarity in the early embryo. We therefore examined PAR-3 localization in *pig-1* single mutants and in combination with *par-1(RNAi)* and a temperature sensitive mutation *par-1(zu310ts)*. This *par-1* allele (referred to hereafter as *par-1(ts))* exhibits a strong *par-1* phenotype at 25°C, (Table S1, see also Methods (Spilker et al., 2009). In wild-type embryos, the anterior PAR-3 domain extends to 51.6% EL (*E*mbryo *L*ength, where 0% is the anterior-most and 100% is the posterior-most cortical domain). We found that in *pig-1* mutants, the anterior PAR-3 domain extended slightly further into the posterior during anaphase, to 57.4% EL, similar to the PAR-3 domain in *par-1(zu310ts)* embryos (55.8% EL) and *par-1(RNAi)* embryos (63.2 % EL) (Figure 1B, C). Examination of the PAR-3 domain in the *pig-1;par-1(ts)* double mutants as well as in *pig-1;par-1(RNAi)* embryos revealed an even greater loss of restriction of the anterior PAR domain (to 71.4 % and 86.1% EL, respectively) (Figure 1B, C). Thus, *pig-1* and *par-1* act redundantly to restrict the size of the anterior PAR-3 domain at the one-cell stage.

### PIG-1 and PAR-1 are required for spindle orientation in the EMS cell

In addition to the enhancement of polarity defects observed at 25°C, *pig-1;par-1(ts)* mutants raised at the semi-permissive temperature of 19.5°C exhibited a high level of embryonic lethality (73%) compared to *pig-1* and *par-1(ts)* single mutants (9% and 4% respectively, Table S1). This is similar to the previously reported enhancement of *par-1(ts)* by *pig-1(RNAi)* (Morton et al., 2012). However, in that study, it was found that P granules localized normally and second division spindle orientation was unaffected in *pig-1(RNAi);par-1(ts*) two-cell embryos (Morton et al., 2012), raising the question of whether the one-cell polarity defects in the double mutant embryos are the cause of the lethality. We therefore set out to determine if other aspects of early embryonic development are affected in *pig-1;par-1(ts)* embryos. Because our analysis of the *par-1(zu310ts)* allele revealed that it is a slow-inactivating temperature-sensitive allele (see Methods), double mutants were grown and imaged at the semi-permissive temperature of 19.5 °C.

Time-lapse microscopy of *pig-1;par-1(ts)* double mutants at 19.5°C showed that asymmetric spindle displacement in the one-cell embryo and P1 spindle orientation at the two-cell stage were normal (N =12), similar to what was reported for *pig-1(RNAi)* in the *par-1(ts)* background (Morton et al., 2012). However, at the four-cell stage, the EMS spindle of the *pig-1;par-1(ts)* double mutants was misoriented. In wild-type embryos, the EMS nuclear-centrosome complex undergoes a rotation prior to nuclear envelope breakdown, so that the spindle forms along the A/P axis. We found that over 50% of *pig-1;par-1(ts)* embryos imaged at 19.5°C exhibited abnormal spindle orientations in the EMS cell (N=32). In comparison, single *pig-1* or *par-1(ts)* mutants imaged at 19.5°C showed only a low level of late spindle alignment or L/R divisions (Figure 2B, N= 17 and 27). In embryos raised and filmed at the non-permissive temperature of 25°C, P1 division orientation was still normal in *pig-1;par-1(ts)* double mutants (N=8); however, the frequency of L/R division defects in EMS was 100%, as seen for *par-1(ts)* alone (N=8, 9). Together these observations reveal that there is redundancy between *par-1* and *pig-1* for EMS division orientation, but not for P1 division orientation.

Since the P2 blastomere signals to EMS to guide its spindle positioning and cell fate specification, we examined *pig-1;par-1(ts)* double mutants for defects in cell fate specification at the four-cell stage. P granules are a germline marker that are segregated to the P1 cell and the P2 cell in a PAR dependent manner during the first divisions of wild-type embryos (Kemphues et al., 1988; Mello et al., 1996; Strome and Wood, 1982). We found that in *pig-1* and *par-1(ts)* single and double mutants raised at 19.5°C, P granules were normally localized at 2- and 4-cell stages (N=4, N=30, N=6, respectively) as in wild-type controls (N=15). PIE-1 is a determinant of germ line fate that is also enriched in the posterior P blastomeres at each of the early divisions. PIE-1 was normally localized in two and four-cell embryos of *pig-1;par-1* double mutants (N=11), as in wild type (N=27), at 19.5°C (Figure 2A). These data suggest that at least some aspects of the P1 asymmetric division and subsequent fate of P2 are preserved in *pig-1;par-1(ts)* double mutants at the semi-permissive temperature.

To directly assay the ability of the MES-1/SRC-1 pathway to activate SRC-1 in the *pig-1;par-1(ts)* double mutants at the semi-permissive temperature, we immunostained embryos with the pY99 a monoclonal antibody that recognizes phosphotyrosine. Previous work showed that wild-type embryos show a relative enrichment of pY99 signal at the P2/EMS interface compared to other cell-cell contacts; the pY99 signal is uniform in *mes-1* mutants and is abolished in *src-1* mutants (Bei et al., 2002). To quantify cortical enrichment of pY99 at the P2/EMS interface, we measured the average pixel intensities at the P2/EMS interface and ABp/EMS interfaces. In wild-type embryos, pY99 staining was enriched approximately 2.3 fold at the P2/EMS contact site compared to the ABp/EMS interface. This enrichment was abolished in *mes-1* mutants (Figure 3A, 3B). In *pig-1* and *par-1(ts)* single mutants, enrichment was also evident at the P2/EMS contact. Although the average enrichment was lower than in wild type, 100% of *par-1(zu310ts)*, and 85% of *pig-1(gm344)* embryos showed pY99 enrichment that fell within the wild-type range, compared to 0% of *mes-1* embryos (Figure 3B). Thus, the majority of single mutant *pig-1* or *par-1(ts)* embryos appear to have intact MES-1/SRC-1 signaling. The enrichment observed in *pig-1;par-1(ts)* double mutants was on average lower than either single mutant; nonetheless, 79% of those embryos had some pY99 enrichment at the P2/EMS contact site compared to the ABap/EMS contact site (cortical ratio greater than 1.0) and the enrichment was within the wild-type range in 58% of embryos (Figure 3B). Together, the P granule, PIE-1 and the pY99 localization data suggest that many aspects of P2 and EMS cell fate are normal in the single *par-1(ts*) and *pig-1* mutants at the semi-permissive temperature. The *pig-1;par-1(ts)* double mutant however, does show a small but signification reduction in pY99 signal between P2 and EMS compared to the single mutants. MES-1 was previously shown to be completely absent in a strong par-1 mutant (Berkowitz and Strome, 2000). Thus, the observations of the *pig-1;par-1* double mutant are consistent with PIG-1 and PAR-1 being partially redundant for proper MES-1/SRC-1 signaling.

**Figure 3.**
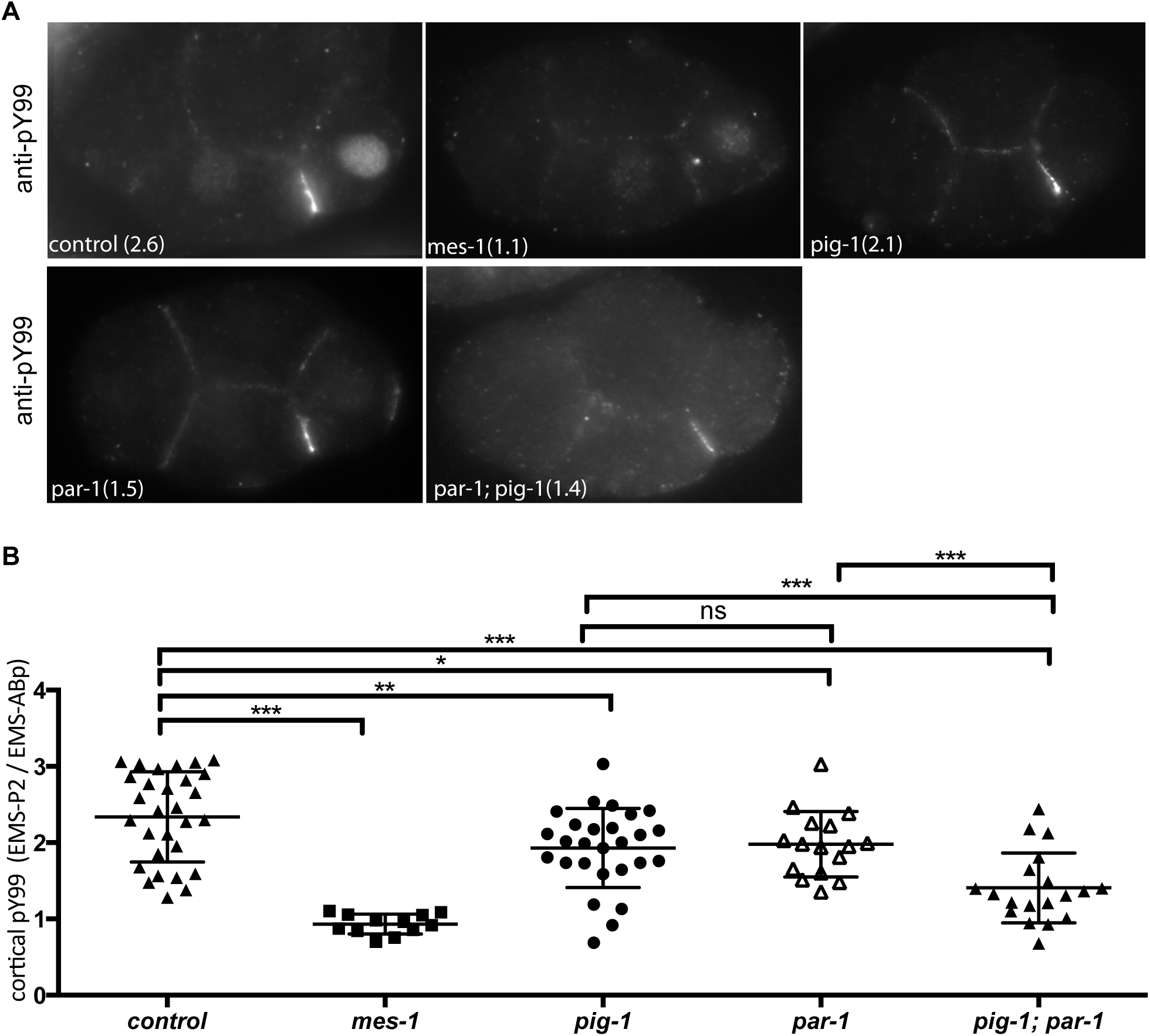
Phospho-tyrosine is enriched at the P2-EMS contact site in *pig-1* and *par-1* mutants. (A) Anti-phospho tyrosine antibody staining in wild type control (N2), *mes-1(bn74)*, *pig-1*(gm344), *par-1(zu310)* and *pig-1(gm344);par-1(zu310)* embryos at 19-20°C. Values in parentheses refer to the ratio of the pY99 signal at the EMS-P2 cortex over the EMS-ABp cortical signal for the specific image shown. (B) Scatterplot of the ratio of pY99 signals at the EMS-P2 cell-cell contact site over EMS-ABp contact site for individual embryos of each genotype indicated. Data were compared using the unpaired Student’s t-test (ns, not significant, P>0.05;* P ≤ 0.05; **P≤0.01; *** P ≤ 0.001; see Table S2 for specific P values).

### PIG-1 acts in the MES-1/SRC-1 pathway for EMS spindle positioning and intestinal specification

The EMS spindle positioning defects observed in the *pig-1* mutant are reminiscent of mutants in the Wnt or MES-1/SRC-1 pathways, which exhibit a low frequency of failed or late EMS spindle rotations (Bei et al., 2002; Liro and Rose, 2016; Schlesinger et al., 1999). Double mutants of components in the same pathway do not show any enhancement of spindle positioning defects as compared to single mutants alone. In contrast, the majority of *wnt;mes-1* double mutants exhibit a complete failure of EMS nuclear/spindle rotation so that the spindle remains on the L/R or other non-A/P axis. We therefore imaged *pig-1* embryos in combination with depletion of Wnt or MES-1/SRC-1 pathway components via RNAi to determine if PIG-1 acts in either pathway. The EMS spindle positioning defects of *pig-1* embryos were enhanced by RNAi of the Wnt components *mom-2* (the ortholog of the vertebrate Wnt ligand) and *dsh-2;mig-5* (vertebrate Dsh orthologs), to 69% and 73%, respectively (Figure 4A). EMS spindle orientation defects seen in *pig-1;wnt(RNAi)* double mutants were comparable to those seen in *mes-1;wnt(RNAi)* embryos. However, *mes-1(RNAi)* did not enhance *pig-1* single mutant EMS spindle positioning defects (Figure 4A). These results are consistent with PIG-1 acting in the MES-1/SRC-1 pathway for spindle orientation.

**Figure 4.**
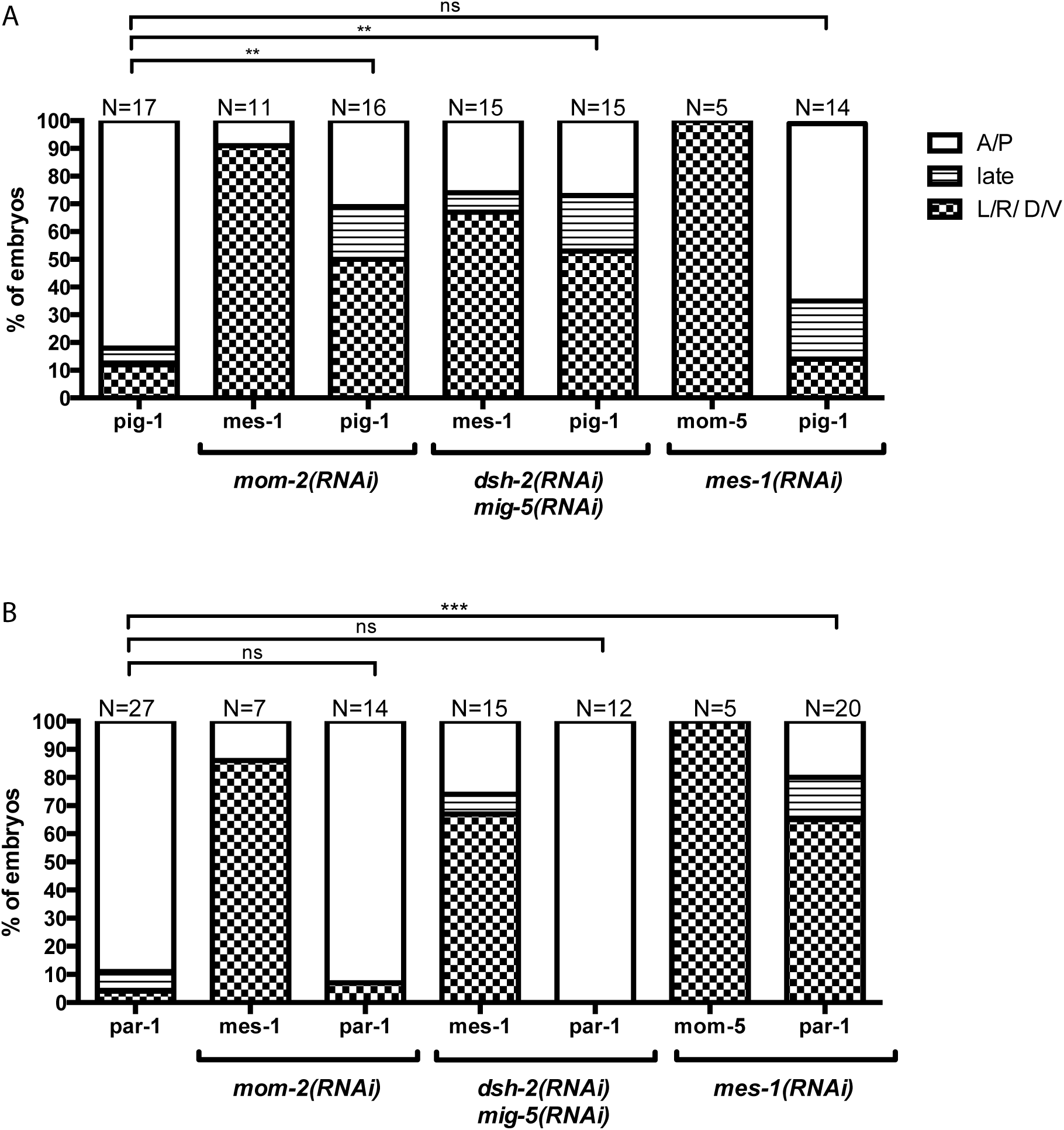
PIG-1 acts in the MES-1/SRC-1 pathway and PAR-1 in the Wnt pathway for EMS spindle positioning. (A) and (B) Quantification of EMS spindle positioning in embryos grown at the semi-permissive temperature of 19.5°C. Spindle orientation was scored as in Figure 2. Single mes-1(RNAi) or wnt(RNAi) treatments were carried out in parallel: *mom-2(RNAi) exhibited* 15/15 A/P divisions, *dsh-2(RNAi); mig-5(RNAi)* embryos exhibited 9/9 A/P divisions, and *mes-1(RNAi)* embryos exhibited 7/8 A/P divisions and 1/8 late rotation. Data were compared using Chi-squared analysis (ns, not significant, P>0.05; * P ≤ 0.05; **P≤0.01; *** P ≤ 0.001; see Table S2 for specific P values).

To determine if PIG-1 also plays a role specifying endoderm fate in the MES-1/SRC-1 pathway, we examined embryos for the presence of gut cells, which are derived from the E daughter of EMS. Single *mes-1* mutants do not have defects in endoderm specification, and many *wnt* mutants display a low frequency of the endoderm specification phenotype. However as with spindle positioning, endoderm defects are enhanced in *wnt;mes-1* double mutants (Bei et al., 2002). Embryos were examined for the presence of autofluorescent gut granules, a marker of gut cells, at a time when control embryos had hatched into larvae (Laufer et al., 1980). As previously reported, only a small fraction of control embryos treated with *mom-2(RNAi),* or *dsh-2(RNAi);mig-5(RNAi)* failed to produce gut granules (Figure 5). While all *pig-1* single mutant late-stage embryos had gut granules, *pig-1; mom-2(RNAi)* and *pig-1;dsh-2(RNAi);mig-5(RNAi)* exhibited a gutless phenotype at high frequency of 57%, which is similar to that seen in *mes-1; mom-2(RNAi)* and *mes-1;dsh-2(RNAi);mig-5(RNAi)* double mutants. In contrast, none of the *pig-1; mes-1(RNAi)* exhibited a gutless phenotype (Figure 5B). Together, these results suggest that PIG-1 acts in the MES-1/SRC-1 pathway for intestinal cell fate specification.

**Figure 5.**
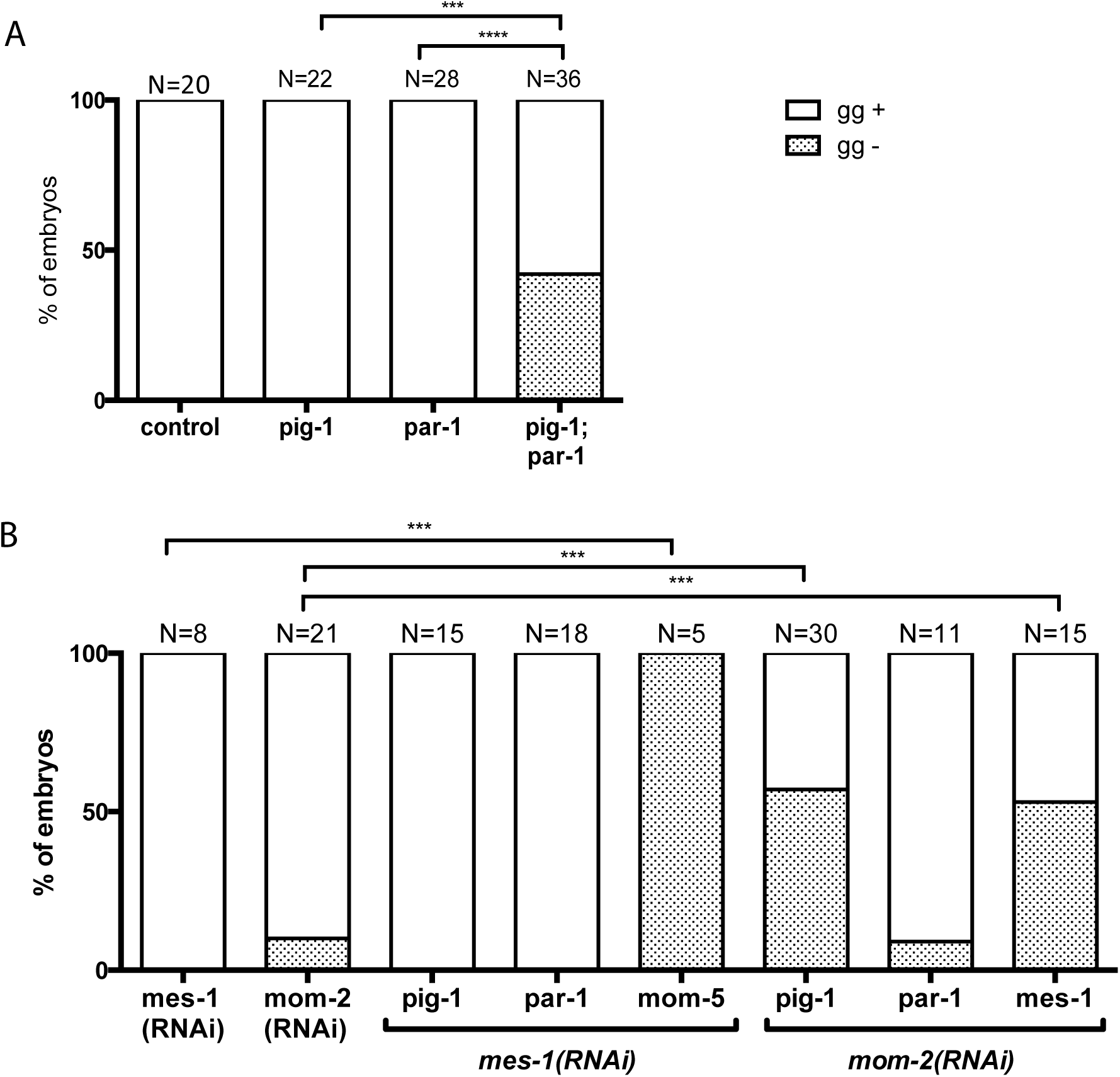
PIG-1 acts in the MES-1 pathway for intestinal specification. Quantification of the presence (gg+) or absence (gg−) of gut granules as a marker for intestinal differentiation. (A) *pig-1(gm344), par-1(zu310)*, *mes-1(RNAi)*, *dsh-2(RNAi);mig-5(RNAi),* and *mom-2(RNAi)* single mutants and (B) in double mutant combinations. Data were compared using Chi-squared analysis. (ns, not significant, P>0.05,* P ≤ 0.05,**P≤0.01, *** P ≤ 0.001, see Table S2 for specific P values).

### PAR-1 acts in the Wnt pathway for EMS spindle positioning

Strong *par -1* loss-of-function mutations show a complete loss of endoderm (Kemphues et al., 1988). At the non-permissive temperature of 25 degrees, *par-1(ts)* single mutants showed a reduction in endoderm specification (80 % gutless embryos, n =35) and 100% L/R spindle orientations (Fig 2B). The gut specification phenotype was enhanced to 100% in pig-1; par-1(ts) double mutant embryos (n= 21). Given PAR-1’s role in early divisions and cytoplasmic asymmetry these defects are likely due at least in part to abnormalities in P2 cell fate that may abrogate both Wnt and MES/SRC-1 signaling.

Because *par-1(zu310ts)* is a slow-inactivating allele, we were unable to perform temperature upshift experiments to test its role more specifically in the EMS division. Instead we again examined *par-1(ts)* at the semi-permissive of 19.5°C, in combination with RNAi of Wnt or MES-1/SRC-1 pathway components. The EMS spindle positioning defects of *par-1(ts)* embryos at 19.5°C were enhanced to 80% when combined with *mes-1*(RNAi), but no enhancement was observed with *mom-2 (RNAi)* (Figure 4B). Neither *wnt* nor *mes-1* RNAi treated *par-1(ts)* embryos showed enhancement of the gutless phenotype at the semi-permissive temperature (Figure 5B). However, *pig-1;par-1(ts)* embryos did exhibit a strong gutless phenotype (Figure 5A), at 19.5°C degrees. These data are consistent with PAR-1 acting in the Wnt pathway, in parallel to the MES-1/SRC-1 pathway, for at least EMS spindle positioning.

## Discussion

Our study provides insight into the role of PIG-1 in regulating the domains of the conserved and well-studied PAR proteins, and also reveals that PIG-1 plays a role in another asymmetric cell division, that of the EMS blastomere.

Multiple mechanisms ensure that polarity is established prior to the first division of the *C. elegans* embryo; PAR proteins need to be localized to distinct anterior vs posterior domains for successful asymmetric division. The first mechanism includes an unknown cue from the sperm centrosome that causes actomyosin flow towards the opposite, the anterior end; this results in concomitant movement of PAR-3, PAR-6 and PKC-3 to the anterior (Munro et al., 2004), and is stabilized by feedback between clustered PAR protein complexes (Sailer et al., 2015). In a secondary mechanism, the microtubules nucleating from the posterior sperm centrosome mediate posterior cortical PAR-2 association, which then recruits PAR-1 enabling it to phosphorylate PKC-3, restricting it to the anterior domain (Motegi et al., 2011; Zonies et al., 2010). Polarization kinetics are slower when one of those mechanisms is compromised. Further, although an anterior PAR domain can be established in the absence of PAR-2, PAR-2 is essential during the maintenance phase to prevent anterior PARs from invading the posterior domain (Rose and Gonczy, 2014; Wu and Griffin, 2017). Previous studies showed a weaker effect on polarity maintenance in *par-1* mutants compared to *par-2* mutants (Cuenca et al., 2003). In addition, *par-1* single mutants still exhibit a pseudocleavage furrow, although its extent is slightly reduced (Kirby et al., 1990) In contrast, our observations revealed that *pig-1* mutants have greatly reduced cortical contractility during polarity establishment. Thus, the failure of *pig-1;par-1* double mutants to restrict the anterior complex could be explained by reduced NMY-2 activity in the *pig-1* mutant, combined with reduced PAR-1 activity in the PAR-2 pathway. Surprisingly however, a recent study concluded that PIG-1 inhibits cortical accumulation of activated myosin, rather than promotes it, during cytokinesis in the *C.elegans* the one-cell (Pacquelet et al., 2015). These different results suggest that PIG-1 may have opposite effects on acotmyosin contractility in the same cell at different time points, likely by regulating different intermediates.

Our data also indicate that PIG-1 is required for both spindle positioning and endoderm fate specification in the EMS cell in the MES-1/SRC-1 pathway. NMY-2 has been placed in the MES-1/SRC-1 pathway for endoderm specification based on enhancement of *wnt* mutants, however spindle orientation was not affected. Further, it was found that *nmy-2(ts)* mutants showed reduced enrichment of the phosphotyrosine marker for SRC-1 activation at the P2/EMS boundary, suggesting that NMY-2 acts upstream of SRC-1 (Liu et al., 2010). In contrast, we found that the enrichment for pY99 was normal in the majority of *pig-1* single mutants, and *pig-1* enhanced both spindle positioning and endoderm specification defects of *wnt(RNAi)* embryos. Thus, PIG-1 likely acts downstream of SRC-1 in the EMS cell to promote asymmetric division. At the same time, we cannot rule out a partially redundant role for PIG-1 in the P2 cell as well, given the reduction in pY99 staining exhibited by *pig-1;par-1(ts)* double mutants at semi-permissive temperature compared to *pig-1* or *par-1(ts)* single mutants. *par-1(ts)* mutant embryos at high temperature as well as other strong *par-1* mutants have a complete loss of endoderm (Kemphues et al., 1988); this could be due to defects in the first or second asymmetric division that result in a failure to localize determinants to the P2 cell, or to a role in generating the MES-1 and Wnt signals at the 4-cell stage. Because the *par-1(ts)* allele is not fast-inactivating, we cannot distinguish between these possibilities.

In addition to specifying general P2 fate, we speculate that PAR-1 may also play a more direct role in the Wnt pathway for EMS division. Our analysis of *par-1(ts)* mutants at the semi-permissive temperature showed that spindle positioning defects were enhanced by MES-1 depletion, but not a Wnt component depletion. Similarly, *pig-1;par-1(ts)* embryos showed an enhancement of spindle positioning defects, consistent with PAR-1 and PIG-1 acting in parallel pathways to promote EMS spindle orientation. Surprisingly, we did not observe an enhancement of endoderm specification defects in *par-1(ts); mes-1* embryos at this temperature. It is possible that spindle positioning is more sensitive to the loss of *par-1* and *mes-1* than is endoderm specification at this temperature. Alternatively, PAR-1 could act downstream of Wnt signaling in the EMS cell to promote spindle positioning. Future studies identifying substrates of PAR-1 and PIG-1 will more fully elucidate their function in EMS cell fate specification and spindle orientation.

In summary, the PIG-1 protein, while nonessential for viability, clearly plays important roles to ensure robust early development in *C. elegans*. Our studies of the parallel requirements of *par-1* and *pig-1* show how important a “nonessential” kinase like PIG-1 is when a more essential effector, such as PAR-1 is weakly compromised. In this situation, roles for PIG-1 in regulating restriction of the anterior polarity protein PAR-3 in the zygote, control of cell division timing in the early embryo, and regulation of EMS spindle orientation and endoderm specification in *C. elegans* were revealed. Our analysis also showed that PAR-1 has a specific role in EMS cell division dynamics in a pathway separable from that of PIG-1. The mammalian homolog of PIG-1, MELK, is also nonessential for viability in mice ((Lin et al., 2017; Wang et al., 2014), but is highly expressed in mammalian embryonic cells, hematopoietic cells and neural progenitor cells (Gil et al., 1997; Heyer et al., 1997; Nakano et al., 2005). MELK is likely to act redundantly with other mitotic kinases, which may include closely related MARK family kinases, homologs of PAR-1 (Drewes et al., 1997). Our results also raise the possibility that mammalian MELK could participate in Src signaling pathways. Further characterization of PIG-1 genetic interactions and substrates in the *C. elegans* system should provide additional insights into the role of this conserved kinase in cell-cell signaling.

## Materials and Methods

### Worm strains and RNAi clones

*C. elegans* were grown using standard conditions (Brenner, 1974). Strains used:

N2: Bristol variant (Brenner, 1974);.

AZ244: unc-119(ed3) III; ruIs57[pie-1::GFP::tubulin + unc-119(+)](Praitis et al., 2001);

KK822: par-1(zu310) V;

KK863: sqt-3(sc8) par-1(zu310) V;

KK1083: pig-1(gm344) IV (derived from NG4370: zdIs5 I; pig-1(gm344) (Cordes et al., 2006);

KK1237: pig-1(gm344) IV; sqt-3(sc8) par-1(zu310) V;

RL262: mom-5(zu193) unc-13(e1091)/hT2 I; +/hT2[bli-4(e937)let(h661)]; unc-119(ed3) III;

ruIs57[pie-1::GFP::tubulin + unc-119(+)] (Liro and Rose, 2016);

RL292: (bn7) X; unc-119(ed3)III; ruIs57[pie-1::GFP::tubulin + unc-119(+)](Liro and Rose, 2016);

RL347: pig-1(gm344) IV; ruIs57[pie-1::GFP::tubulin + unc-119(+)];

WM150 pkc-3(ne4246ts) and WM151 pkc-3(ne4250ts) (Fievet et al., 2013);

mes-1(bn74) X (Berkowitz and Strome, 2000).

Temperature-sensitive strains were maintained at 16.0°C +/-1.0°C and worms were shifted to 19.5 +/-0.5°C or 25 +/-0.5°C as L3/L4 for all analyses. Other strains were maintained at 19.5 °C +/-1.0°C.

RNAi was carried out by bacterial feeding (Timmons and Fire, 1998) using the following Ahringer library clones: mes-1(X-5L23), mom-2(V-6A13), mig-5(II-6C13), dsh-2(II-4011) (Kamath et al., 2003), and a *par-1* RNAi clone (Hurd and Kemphues, 2003). Worms were imaged 20-30hr post shift to 19.5°C.

### Live imaging

Embryos were removed from gravid hermaphrodites and were mounted on 2% agarose pads on microscope slides and covered with coverslip. The initial time-lapse microscopy of *pig-1(gm344), par-1(zu310)* and the double *pig-1(gm344);par-1(zu310)* mutant embryos was carried out using DIC optics on a Leica DM RA2 microscope with a Hamamatsu ORCAER digital camera or on an Olympus BX60 fitted with an Hamamatsu Orca C4742-95 camera, using Openlab or ImagePro software in a temperature-controlled room kept at (19.5°C). Subsequent analyses including combinations with RNAi were carried out using an Olympus BX60 microscope a Hammatasu Orca 12-bit digital camera fitted with a Linkam PE95/T95 System Controller with an Eheim Water Circulation Pump set to maintain the temperature of the slide at 19 +/-0.5 °C or 25 +/-0.5 °C (Settings were 15°C and 25°, true temperatures were determined by inserting the wire probe of an Omega HH81 digital thermometer between the cover slip and an agar pad with the 60X objective and oil in place). Single-plane images in brightfield optics were acquired every 10 sec using Micromanager 1.4.22. EMS spindle positions were categorized as in Liro and Rose (2016): A/P, spindle initially formed on the A/P axis; late, spindle formed on the L/R or other non-A/P axis and then rotated onto the A/P axis; L/R D/V, the spindle formed on a non-A/P axis and never rotated. Phenotypes were grouped as normal (A/P) or abnormal (late or L/R D/V) and compared using Chi-squared analysis in Excel. Graphs were made in GraphPad Prism Version 6.0.

*par-1(zu310)* was found to be a slow-inactivating temperature sensitive allele. Embryos from worms shifted from 16°C to 25°C for 18-24 hr showed synchronous cleavages and 100% L/R orientation of the EMS cell division (N=9). In contrast, embryos shifted to 25°C for 10 min up to 100min (N=8) appeared the same as those grown at 16, with more asynchronous divisions and normal EMS division orientations.

### Immunofluorescence and quantification of staining

All embryos used for PAR-3 staining were from worms grown at 25°C, while embryos stained with other antibodies were from worms grown at 19.5°C from the L3/L4 stage. Worms were dissected in water of the same temperature on slides, frozen in liquid nitrogen, then fixed in methanol at -20°C for 15 min. For P granule staining, a post methanol acetone step was included (Strome and Wood, 1983). Primary antibodies used were monoclonal mouse anti–PAR-3 (Nance et al., 2003), OIC1D4 monoclonal mouse anti-P granule (Strome, 1986), anti-PIE-1 (Tenenhaus et al., 1998) and pY99 (Santa Cruz Biotechnology). Monoclonal antibodies were obtained from the Developmental Studies Hybridoma Bank, University of Iowa. Secondary antibodies included Alexa Fluor 594-labeled goat anti-mouse (Invitrogen) and Cy3-labeled donkey anti-mouse (Jackson ImmunoResearch). Slides were mounted with Vectashield containing DAPI (Vector Laboratories). Images were acquired on a Leica DM RA2 microscope fitted with a Hamamatsu Orca-ER digital Camera using OpenLab software (Improvision).

OpenLab software was used to measure length of PAR-3 anterior–posterior domain and overall length of embryos in pixels, which was then converted to percent embryo length. Measurements were taken twice for each embryo. For quantification of pY99 staining, four-cell embryos where P2 and EMS were in interphase to metaphase 4-cell embryos were used. Fluorescence intensities were traced along the EMS-P2 and EMS-ABp cell-cell contacts using the segmented line tool of the Fiji software. The ratios of EMS-P2 cortical to the EMS-ABp cortical pixel intensities were then calculated for each embryo, and were averaged for each genotype. Statistical tests of significance were made using the Student’s t-test in Excel and the scatterplot was created in GraphPad Prism Version 6.0.

### Analysis of gut cell fate specification

Gut cell differentiation was scored in the same embryos filmed for the spindle positioning data or in embryos from siblings worms raised in parallel, that were mounted on agar pads on slides. Slides were incubated in a moist chamber until the time that normal embryos would hatch (at least 12 hours at 25°C or 24 hours at 19.5 °C). UV light or polarization optics were used to identify the presence autofluorescent/birefringent gut granules (Laufer et al., 1980).

## Acknowledgements

We thank Gian Garriga, Craig Mello and Susan Strome for strains and Jim Priess’s lab for PIE-1 antibody. Some strains were provided by the CGC, which is funded by NIH Office of Research Infrastructure Programs (P40 OD010440). We are grateful to Neil Willits, UC Davis Department of Statistics, for assistance with the Chi-squared analysis. We thank members of the Rose and McNally labs, the UC Davis SuperWorm group, Ken Kemphues, members of the Kemphues lab, and Kelly Liu for helpful discussion. We also thank Ken Kemphues for helpful comments on the manuscript. This work was supported by NIH 1R01GM68744 and by NIFA CA-D* -MCB-6239-H to L.R. and by NIH R01GM079112 to Kenneth Kemphues.

---

**Table S1.**
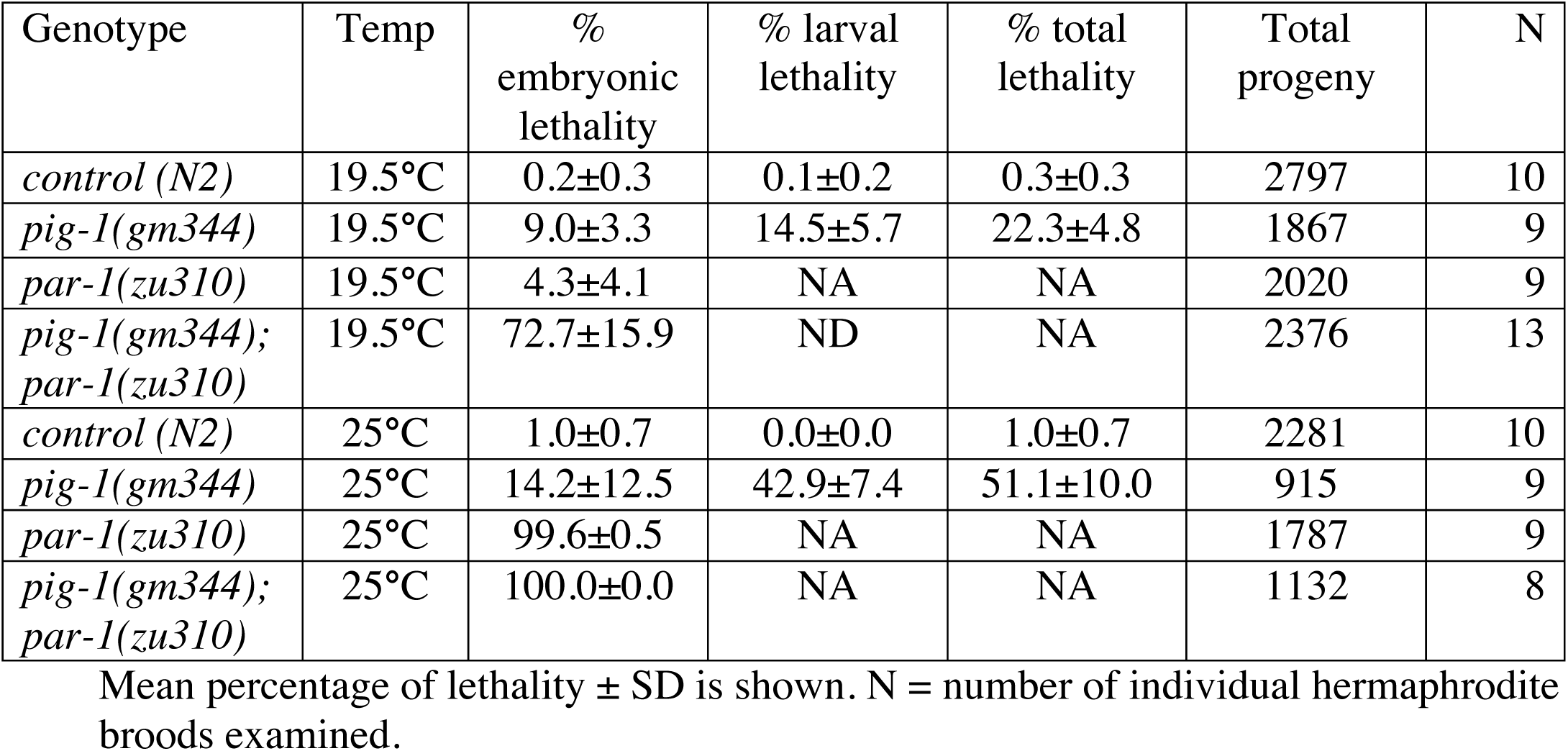
Lethality of *pig-1* single mutants compared to *par-1* single and *pig-1;par-1* double mutants.

**Table S2.** Statistical comparisons

| **Figure** | **Genotypes compared** | **P value** |  |
| --- | --- | --- | --- |
| **Figure 1C** | *control vs pig-1* | 0.0061 | ** |
|  | *control vs par-1* | 0.016 | * |
|  | *control vs pig-1;par-1* | <<0.0001 | **** |
|  | *pig-1 vs par-1* | 0.41 | ns |
|  | *pig-1 vs pig-1;par-1* | <<0.0001 | **** |
|  | *par-1 vs pig-1;par-1* | <<0.0001 | **** |
|  | *par-1(RNAi) vs pig-1;par-1(RNAi)* | <<0.0001 | **** |
| **Figure 2A** | *Control vs pig-1 ($\Delta$ AB-P1)* | 0.00030 | *** |
|  | *Control vs pig-1 ($\Delta$ P2-EMS)* | 0.16 | ns |
|  | *par-1 vs pig-1;par-1 ($\Delta$ AB-P1)* | 0.133 | ns |
|  | *par-1 vs pig-1;par-1 ($\Delta$ P2-EMS)* | 0.055 | ns |
| **Figure 2B** | *Control vs pig-1 20°C* | 0.050 | ns |
|  | *Control vs par-1 20°C* | 0.12 | ns |
|  | *pig-1 vs pig-1;par-1 20°C* | 0.0093 | ** |
|  | *par-1 vs pig-1;par-1 20°C* | 0.00031 | *** |
|  | *pig-1 vs par-1 25°C* | 0.0025 | ** |
|  | *pig-1 vs pig-1;par-1 25°C* | 0.0016 | ** |
| **Figure 3B** | *control vs mes-1* | <<0.0001 | **** |
|  | *control vs pig-1* | 0.00834 | ** |
|  | *control vs par-1* | 0.039 | * |
|  | *control vs pig-1;par-1* | <<0.0001 | **** |
|  | *pig-1 vs mes-1* | <<0.0001 | **** |
|  | *pig-1 vs par-1* | 0.748 | ns |
|  | *pig-1 vs pig-1;par-1* | 0.0010 | ** |
|  | *par-1 vs mes-1* | <<0.0001 | **** |
|  | *par-1 vs pig-1;par-1* | 0.000614 | *** |
|  | *mes-1 vs pig-1;par-1* | 0.0016 | ** |
| **Figure 4** |  |  |  |
|  | *pig-1 vs pig-1;mom-2(RNAi)* | 0.0030 | ** |
|  | *mom-2(RNAi) vs pig-1;mom-2(RNAi)* | 0.00006 | **** |
|  | *pig-1 vs pig-1; dsh-2(RNAi); mig-5(RNAi)* | 0.00153 | ** |
|  | *dsh-2(RNAi); mig-5(RNAi) vs pig-1; dsh-2(RNAi); mig-5(RNAi)* | 0.00048 | *** |
|  | *pig-1 vs pig-1;mes-1(RNAi)* | 0.25 | ns |
|  | *mes-1(RNAi) vs pig-1;mes-1(RNAi)* | 0.24 | ns |
|  | *par-1 vs par-1;mom-2(RNAi)* | 0.68 | ns |
|  | *mom-2(RNAi) vs par-1;mom-2(RNAi)* | 0.29 | ns |
|  | *par-1* vs *par-1; dsh-2(RNAi); mig-5(RNAi)* | 0.23 | ns |
|  | *par-1* vs *par-1;mes-1(RNAi)* | <0.0001 | **** |
|  | *mes-1(RNAi)* vs *par-1;mes-1(RNAi)* | 0.00095 | *** |
| **Figure 5** |  |  |  |
|  | *pig-1* vs *pig-1;par-1* | 0.00044 | *** |
|  | *par-1* vs *pig-1;par-1* | 0.000090 | **** |
|  | *mom-2(RNAi)* vs *pig-1; mom-2(RNAi)* | 0.00061 | *** |
|  | *mom-2(RNAi)* vs *par-1; mom-2(RNAi)* | 0.97 | ns |
|  | *mes-1(RNAi)* vs *par-1; mes-1(RNAi)* | 0.50 | ns |

